# Sarcomere dynamic instability and stochastic heterogeneity drive robust cardiomyocyte contraction

**DOI:** 10.1101/2024.05.28.596183

**Authors:** Daniel Haertter, Lara Hauke, Til Driehorst, Kengo Nishi, Wolfram-Hubertus Zimmermann, Christoph F. Schmidt

**Affiliations:** Institute of Pharmacology and Toxicology, University Medical Center Göttingen, Göttingen, Germany; DZHK (German Center for Cardiovascular Research), partner site Göttingen, Göttingen, Germany; CIDAS (Campus Institute Data Science), University of Göttingen, Göttingen, Germany; Department of Physics and Soft Matter Center, Duke University, Durham, NC, USA; Third Institute of Physics, Faculty for Physics, University of Göttingen, Göttingen, Germany; Cluster of Excellence “Multiscale Bioimaging: from Molecular Machines to Networks of Excitable Cells” (MBExC), University of Göttingen, Germany; Fraunhofer Institute for Translational Medicine and Pharmacology, Göttingen, Germany

## Abstract

Cardiac contraction is driven by the collective action of cardiomyocytes, that contain parallel bundles of myofibrils consisting of linear chains of sarcomeres, the basic force-generating units. The dynamics of individual sarcomeres within intact cardiomyocytes remain incompletely understood. While most models assume uniform, synchronized contractions, recent studies hint at unexpected heterogeneity whose origins and significance are not yet clear. By combining the culture of fluorescent sarcomere-reporter hiPSC-derived cardiomyocytes on micropatterned soft gels of different stiffness (5 – 85 kPa) with AI-based tracking of sarcomere motion, we found that increasingly stiff substrates inhibited overall cardiomyocyte contraction, but, surprisingly, did not diminish individual sarcomere dynamics. Instead, sarcomeres competed in a tug-of-war causing increasing heterogeneity, including rapid length oscillations and overextensions (popping). Statistical analysis showed that the heterogeneous dynamics were not caused by static structural differences but were largely stochastic. Stochastic heterogeneity is thus an intrinsic property of cardiac sarcomeres and likely mediates the adaptation of cardiomyocyte contractility to mechanical constraints. A mesoscopic model of coupled sarcomeres shows that these phenomena can be explained by a non-monotonic force-velocity relationship and stochastic fluctuations, where dynamic instability at a critical yielding force creates heterogeneity. Stochastic heterogeneity compensates for structural disorder by randomizing yield events beat-to-beat, preventing damage to specific sarcomeres. Our findings recast cardiac sarcomeres as active, dynamically unstable, and stochastic units engaged in a stochastic tug-of-war, where transient, velocity-dependent forces dominate. We propose that pathological disorder in cardiomyopathy drives a transition from protective stochastic fluctuations to more deterministic, persistently overloaded sarcomeres.

## Introduction

Sarcomeres are the basic contractile cytoskeletal units of striated muscle, including cardiac muscle. In most species, sarcomeres are ∼2 µm in length and consist of highly ordered bipolar bundles of myosin motor proteins interdigitated with actin filaments, tied together in the Z-bands. (**Fig. 1A**). Force generation by myosin interacting with actin is regulated by intracellular Ca^2+^ and fueled by chemical energy (ATP)^1^. Sarcomeres are serially connected into myofibrils that span the length of the cell. A cardiomyocyte (CM) contains tens of parallel myofibrils. Overall CM contractility is a mesoscopic process that emerges from a very large number of coupled, non-linear, stochastic, elementary force generators. Despite extensive knowledge of single-molecule motor kinetics^2,3^ and whole-muscle mechanics^4,5^, this intermediate mesoscopic scale—where tens of sarcomeres interact within a myofibril—remains largely unexplored in cardiac muscle. Sarcomeres are mechanically coupled in series, yet each is also an active, excitable unit undergoing rapid, transient activation-relaxation cycles. Do these coupled sarcomeres maintain synchronized motion, or do stochastic fluctuations and non-equilibrium forces drive heterogeneous behavior?

**Figure 1:**
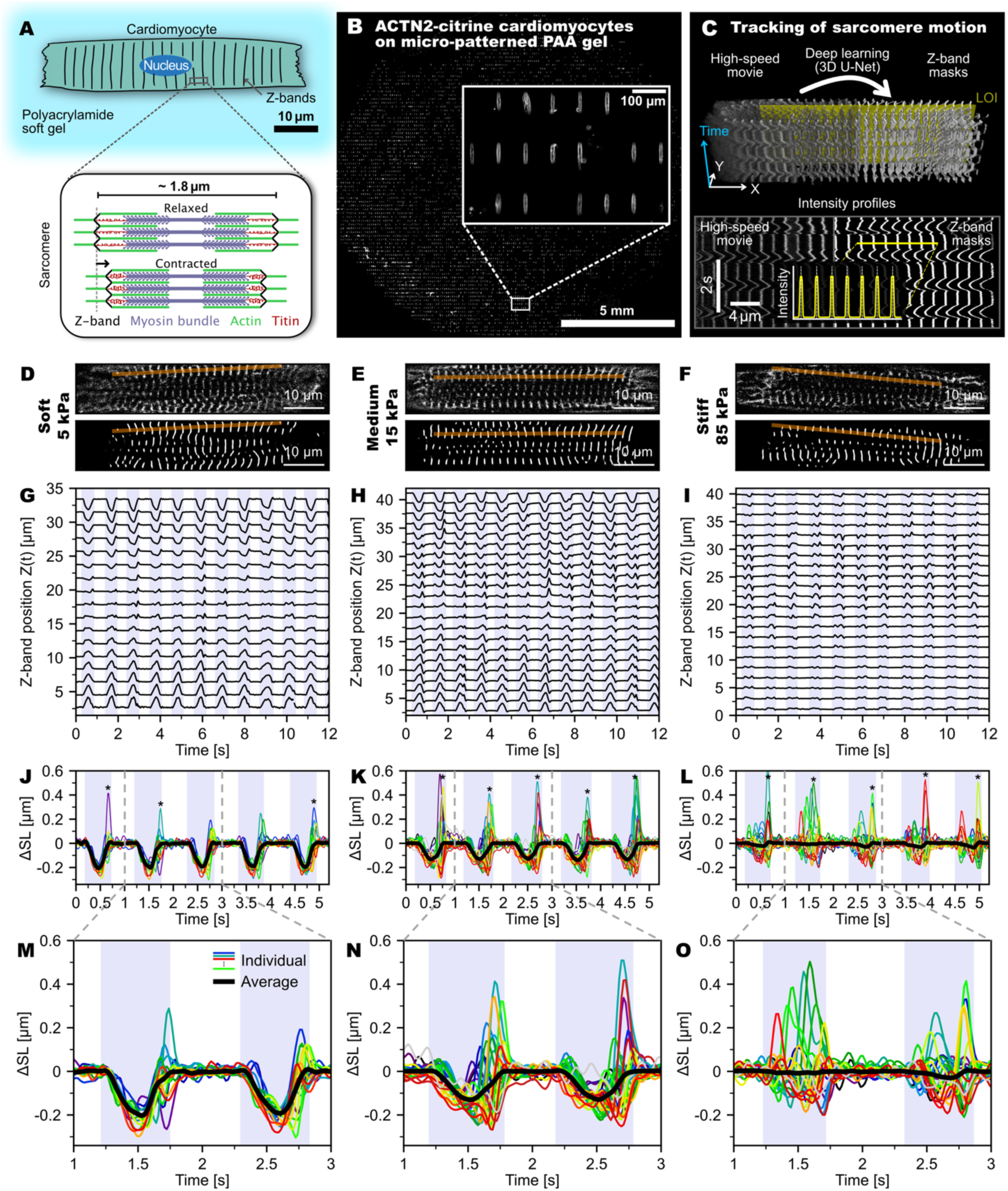
Sarcomere tracking in genetically engineered Z-line labeled iPSC-derived cardiomyocytes on micropatterned soft gels. (**A**) Sketch of a human cardiomyocyte (CM) adhering to a micro-patterned gel (top) and sarcomere structure in relaxed and contracted state (bottom). (**B)** ACTN2-Citrine cardiomyocytes (culture day 20) on a polyacrylamide gel substrate (Young’s modulus: 15 kPa), patterned with rectangular stripes of Synthemax (70 × 10 µm). More than 50% of the stripes were typically occupied by single cardiomyocytes. Inset: zoomed-in view. (**C**) Tracking of sarcomere motion: (Top) blend of high-speed confocal movie of a CM adherent to a 15 kPa substrate (left) and deep-learning (3D U-Net)-based segmentation of sarcomere Z-bands (right) with line of interest (yellow). (Bottom) Intensity extracted along line of interest (LOI). Inset shows representative section of the Z-band intensity profile. (**D**-**E**) Confocal images of representative ACTN2-Citrine-labeled CMs with corresponding Z-band segmentation and LOIs (red lines). (**G-I**) Z-band trajectories tracked along the LOI. (**J-O**) Overlay plots of single sarcomere length change Δ*SL*(*t*) for all tracked sarcomeres in the marked LOIs (first 5 seconds in **J-L**, zoomed in **M-O**). Colored lines are individual sarcomere length changes; black lines display average length changes. Contraction intervals are marked by a blue background. Sarcomere popping events are marked with asterisks. Conditions (substrate stiffness): 5 kPa (**D,G,J,M**); 15 kPa (physiological): (**E,H,K,N**); 85 kPa (**F,I,L,O**).

Recent studies, utilizing in vivo imaging and advanced in vitro models, have begun to reveal notable heterogeneity and beat-to-beat variability at the sarcomere level — even as the average myofibril motion remains smooth and regular^6–8^. However, these observations have been limited to murine systems (≈5-8 Hz beating rate vs. ∼1 Hz in humans). While sarcomere non-uniformity has a long history in skeletal muscle research^9–12^, prevailing models of cardiac contraction have historically assumed that sarcomeres behave as uniform, synchronized units, with force generation determined primarily by the steady-state force–length relationship—an idea rooted in the Frank-Starling mechanism and length-dependent activation^4,13^.

Existing attempts to explain cardiac sarcomere heterogeneity remain largely qualitative, invoking static structural differences between sarcomeres—e.g., in strength or resting length—especially pronounced in cardiomyopathy^6–8,14^. Such models propose deterministic mechanisms: ‘weak’ sarcomeres are consistently stretched by ‘strong’ neighbors^14^, diastolic length variations dictate subsequent systolic interactions and potential role-switching between sarcomeres^6,8^, or inherent length variability enables recruitment of more contracting sarcomeres upon cell lengthening through length-dependent activation^7^.

Crucially, these deterministic, primarily length-centric views neglect velocity-dependent active force generation, friction effects, and non-linearities. By assuming a continuous balance between passive and active forces dictated by steady-state relationships, they ignore the non-equilibrium nature of the beating heart, where adaptation kinetics and viscous drag are critical. Consequently, unraveling these dynamics requires novel approaches that resolve the actual motion of individual sarcomeres within their native physiological setting—inside intact, beating cardiomyocytes—tightly integrated with quantitative modeling.

In this study, we cultured human fluorescent sarcomere-reporter hiPSC-derived cardiomyocytes (ACTN2-Citrine-CMs) on micropatterned gel substrates of different stiffnesses to investigate sarcomere dynamics under different mechanical boundary conditions. High-speed imaging, combined with AI-based analysis using our SarcAsM algorithm^15^, enabled us to track average and individual sarcomere motions within myofibrils with nanometer resolution.

Our analysis found strikingly complex dynamic behaviors, including rapid length oscillations, transient ‘popping’ events, and heterogeneous contraction patterns that varied both temporally and spatially along one myofibril. We show that this stochastic heterogeneity appears to arise from competitive, tug-of-war-like interactions among mechanically coupled sarcomeres within the myofibrils. To rationalize these observations, we developed a data-driven mesoscopic model which demonstrates that the emergent heterogeneity at the sarcomere-level can be explained by stochastic fluctuations in conjunction with a non-monotonic force-velocity relation with a critical force threshold. These insights into stochasticity and dynamic instability at the sarcomere level challenge traditional paradigms of myocardial contraction.

## Results

### Individual ACTN2-Citrine hiPSC-derived CMs on micro-patterned elastic substrates

To resolve sarcomere dynamics with high spatial and temporal resolution in PSC-derived cardiomyocytes, we used a CRISPR-engineered induced pluripotent (iPSC) stem cell line, expressing a yellow fluorescing protein as ACTN2-Citrine N-terminal fusion protein after cardiomyocyte differentiation^15^. Seeding of differentiated cardiomyocytes on soft elastic polyacrylamide gels functionalized with a micro-printed pattern (70 × 10 µm; **Fig. 1B**) promoted anisotropic myofibril assembly in uniformly elongated cardiomyocytes. Culturing cells on gels with defined elasticities (Young’s moduli: 5 kPa, 9 kPa, 15 kPa, 29 kPa, 49 kPa, 85 kPa; **Table S1**) allowed us to impose auxotonic loads on the cells on a scale considered to be relevant *in vivo* under physiological (10-20 kPa) and pathological (e.g., fibrosis; ≥30 kPa) conditions^15^. After a maturation period of 20-35 days, we recorded 20-30 s long movies of, in total, 1,362 spontaneously beating CMs at a frame rate of 66 Hz (**Movie S1, Fig. S1**).

### Automated tracking of sarcomere motion at high spatial and temporal resolution

We used SarcAsM, an AI-based software tool for automated segmentation and analysis of sarcomere structure and dynamics^15^ for data evaluation. SarcAsM automatically identified 5,085 lines of interest (LOIs), up to 4 per cell, with well-organized registered sarcomeres (≥10 sarcomeres; **Fig. 1C, S1**). Along each LOI (∼800 nm in width), an intensity kymograph of Z-band motion was extracted from deep-learning (3D U-Net) processed movies, allowing for robust and time-consistent segmentation of Z-bands in noisy high-speed movies (**Fig. 1C**). This approach provided more accurate and robust localization and tracking of Z-band trajectories 𝑍*_i_*(𝑡) of individual sarcomere𝑠 than using raw microscopy data (∼17 nm and 15 ms resolution, details see SarcAsM^15^). From 𝑍*_i_*(𝑡), we obtained sarcomere length 𝑆𝐿*_i_*(𝑡), sarcomere length change Δ𝑆𝐿*_i_*(𝑡) and sarcomere velocity 𝑉*_i_*(𝑡) of each sarcomere 𝑖 as well as multi-sarcomere averages 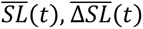 and 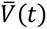 for each LOI (details see SarcAsM^15^) (**Fig. 1D-F**).

### Cardiomyocyte and individual sarcomere contractility as a function of substrate stiffness

The hiPSC-CMs were beating spontaneously at a frequency of 0.91 ± 0.38 Hz, which was largely independent of the substrate stiffness (**Fig. S2A**). Beat-to-beat intervals, though, showed increasing irregularities with increasing substrate stiffness (**Fig. S2B**). Contraction durations 𝑇*_c_* shortened with increasing substrate stiffness (**Fig. S2C**).

In line with previous reports^16–18^, total cell contraction amplitudes, quantified by the inward motion of the outermost z-bands in myofibrils, substantially decreased with increasing substrate stiffness (**Fig. 1D-F**). A surprising finding was that the length changes of individual sarcomeres Δ𝑆𝐿*_i_* fell out of synchronization and became distinctly heterogenous, deviating strongly from the average length change 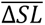, with increasing substrate stiffness. Sarcomeres also showed more singular large-amplitude extensions (“sarcomere popping”) far beyond their resting lengths, often very pronounced at the end of contractions (marked with asterisks in bottom row of **Fig. 1D-F** and large excursions in phase-space plots in **Fig. 2A-C**). Despite the rapid and heterogenous motion of individual sarcomeres, the emergent contraction at the myofibril scale remained regular and smooth on all substrates.

**Figure 2:**
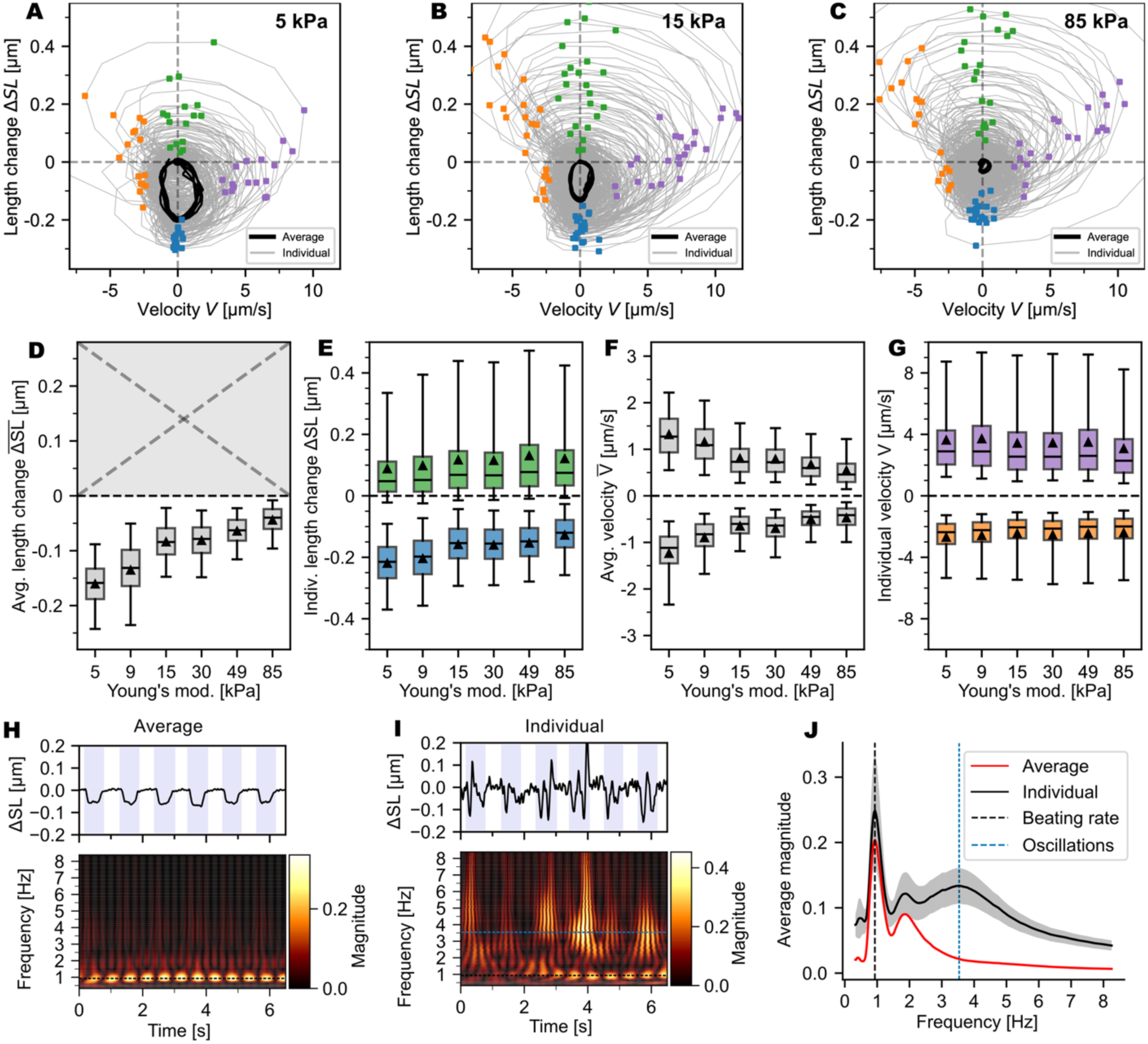
Analysis of sarcomere length change dynamics. (**A-C**) Phase-space plots of sarcomere length change Δ*SL* vs. velocity *V* for three representative LOIs from CMs on substrates of increasing stiffness (same LOIs as in Fig. 1). Gray lines show individual sarcomere dynamics, black lines average dynamics. Colored dots mark maximal and minimal values of Δ*SL* and *V* for individual sarcomere trajectories, with colors corresponding to data in **E** and **G.** (**D**) Box plot of maximal average contractions 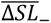 as function of substrate stiffness. Maximal average extensions 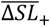 are always close to 0 and not shown. (**E**) Box plot of maximal shortening and lengthening amplitudes Δ𝑆𝐿_+/-_of individual sarcomeres quantified in each contraction cycle. (**F**) Box plot of maximal average sarcomere lengthening and shortening velocities 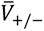. (**G**) Box plot of maximal individual sarcomere lengthening and shortening velocities 𝑉_+/-_. Boxes show quartiles, lines the median, triangles the mean and whiskers the 5th and 95th percentile of the distribution per condition. Each data point corresponds to the extremal value within one contraction cycle. To weigh each LOI equally, only the first 10 contraction cycles in each recording were considered. **D-G** combine data of 1,652 LOIs (5 kPa: 188, 9 kPa: 586, 15 kPa: 394, 29 kPa: 319, 49 kPa: 506, 85 kPa: 347). Statistical analysis was performed using the Kruskal-Wallis and Dunn’s post hoc tests, with significance set at *p* < 0.05. All differences were significant. (**H,I**) Time series and Morlet wavelet scalogram of average and single sarcomere length changes Δ𝑆𝐿(𝑡). The top plot displays average (**H**) and representative single (**I**) sarcomere length changes over time, with blue background marking contraction periods. The bottom plot presents the wavelet scalograms, depicting the evolution of frequency content in the signal over time, with the blue dashed line signifying the cell’s beating rate. (**J**) Comparison of time-averaged oscillation frequencies of average (red) and single (black) sarcomere length changes of one representative LOI, showing high-frequency intrinsic oscillatory motion of individual sarcomeres with frequencies of 3-4 Hz, which cancel out on the myofibril scale. For time-averaging of the frequency spectra, only the contraction intervals were included. The black curve shows mean ± S.D. of 16 sarcomeres in one representative LOI; the black dashed line shows the beating rate and the blue dashed line the peak of the oscillation frequency distribution.

To analyze the deviations of individual sarcomere dynamics from the myofibril average during contraction cycles, we collected minima and maxima of Δ*SL*, i.e., maximal contraction and elongation amplitudes, and minima and maxima of *V* for each contraction cycle for both individual sarcomeres and averages over all sarcomeres in a given LOI (**Fig. 2A-G**). In total, we analyzed a data set of 5,085 LOIs recorded from 1,362 cells, each over at least 10 contraction cycles. We excluded irregularly beating (beat-to-beat variability >0.1 s) and very rapidly (>2 Hz) or slowly (<0.5 Hz) beating cells. We further limited the analysis to LOIs spanning whole myofibrils, where the average length change was strictly negative, i.e., showing contraction. Out of the full data set of 5,085 LOIs, 2,321 LOIs met these criteria. The majority of eliminated LOIs were from irregularly beating cells (**Fig. S2B**) This selection ensured that we focused on rhythmically and uniformly contracting cells. As expected, maximal average contraction amplitudes 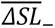, measured at the peaks of shortening in each contraction cycle, were the largest (0.16 ± 0.05 µm) on the softest (5 kPa) substrates and were close to zero (0.04 ± 0.03 µm) on the hardest substrates (85 kPa; **Fig. 2D**). Maximal contraction amplitudes 𝛥*SL*_-_ of individual sarcomeres in each contraction cycle were also largest on 5 kPa substrates at 0.22 ± 0.09 µm and declined to 0.13 ± 0.07 µm on 85 kPa (**Fig. 2E**). Note that the decline of maximal individual sarcomere shortening (𝛥*SL*_-_) by ∼30% from 5 to 85 kPa was far less than the ∼75% decline of the average sarcomere contraction (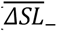). In addition, while myofibrils never elongated beyond resting length, as expected for auxotonic contractions, individual sarcomeres frequently elongated well beyond their resting lengths during contraction cycles (median 𝛥*SL*_+_ = 0.05 µm). The distributions of maximal sarcomere extension amplitudes (𝛥*SL*_+_) were remarkably similar between substrate conditions (**Fig. 2E**).

Next, we evaluated the effect of substrate elasticity on average and individual sarcomere contraction and extension velocities. The average extremal velocities 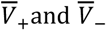 followed the same trend as the extremal average length changes 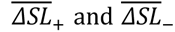 and were the largest on 5 kPa and declined steadily and strongly with increasing substrate stiffness (from 5 to 85 kPa by ∼60% for both contraction and elongation; **Fig. 2F**). The single sarcomere extremal contraction and extension velocities 𝑉_-_and 𝑉_+_, in contrast, were far less affected by substrate elasticity and differed only by at most ∼10% (𝑉_-_) and ∼20% (𝑉_+_) for the various substrate conditions (**Fig. 2G**). Note that the maximal extension velocities were 29 - 38% larger than the maximal absolute contraction velocities, showing a strong asymmetry between single sarcomere contraction and extension dynamics.

### High-frequency oscillatory motion of individual sarcomeres

Unlike the average sarcomere motion with clearly distinguishable contraction and relaxation phases, single sarcomeres exhibited rapid switching between slow contractile and fast extensile motions with sometimes multiple shortening and lengthening phases within one contraction cycle (**Fig, 2H,I, Fig. 1D-F**). To quantify this phenomenon, we analyzed the oscillation frequencies of the average and individual length changes 𝛥𝑆𝐿(𝑡) over time, utilizing Morlet wavelet analysis, a method that makes it possible to extract the instantaneous oscillation frequencies of a signal. For average sarcomere length changes 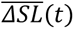, the analysis revealed a narrow range of frequencies surrounding the beating rate in the scalogram (Fig. **2H,I**). For individual sarcomeres, in contrast, we found a broader range of frequencies, with significant peaks occurring at the dominant and macroscopically visible whole-cell beating rate as well as a large secondary peak at 3 ± 0.5 Hz (**Fig. 2J**). This distinctive high-frequency oscillation was consistently observed across numerous LOIs and appeared to be independent of the beating rate. The intrinsic oscillatory motion of individual sarcomeres suggests a mechanism of active acto-myosin powered contraction coupled to rapid lengthening in a relaxation oscillator-like behavior ^19^.

### Correlation analysis identifies stochastic and static heterogeneity among sarcomeres

The observed heterogeneity of sarcomere dynamics, particularly on rigid substrates (**Fig. 3A**), might be predetermined by static non-uniformities among sarcomeres (e.g., by differences in functional myosin numbers). To distinguish this possibility from stochastic heterogeneity, we extracted the time-resolved motion Δ𝑆𝐿*_i,k_*(𝑡) of each sarcomere *i* during each contraction cycle *k*. We then quantified the similarity between these motion patterns using the Pearson correlation coefficient. For two motion patterns Δ𝑆𝐿*_i,k_*(𝑡) and Δ𝑆𝐿*_j,k_*(𝑡), this coefficient is defined as:

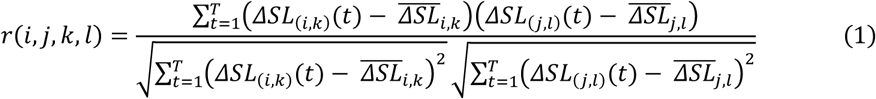

**Figure 3:**
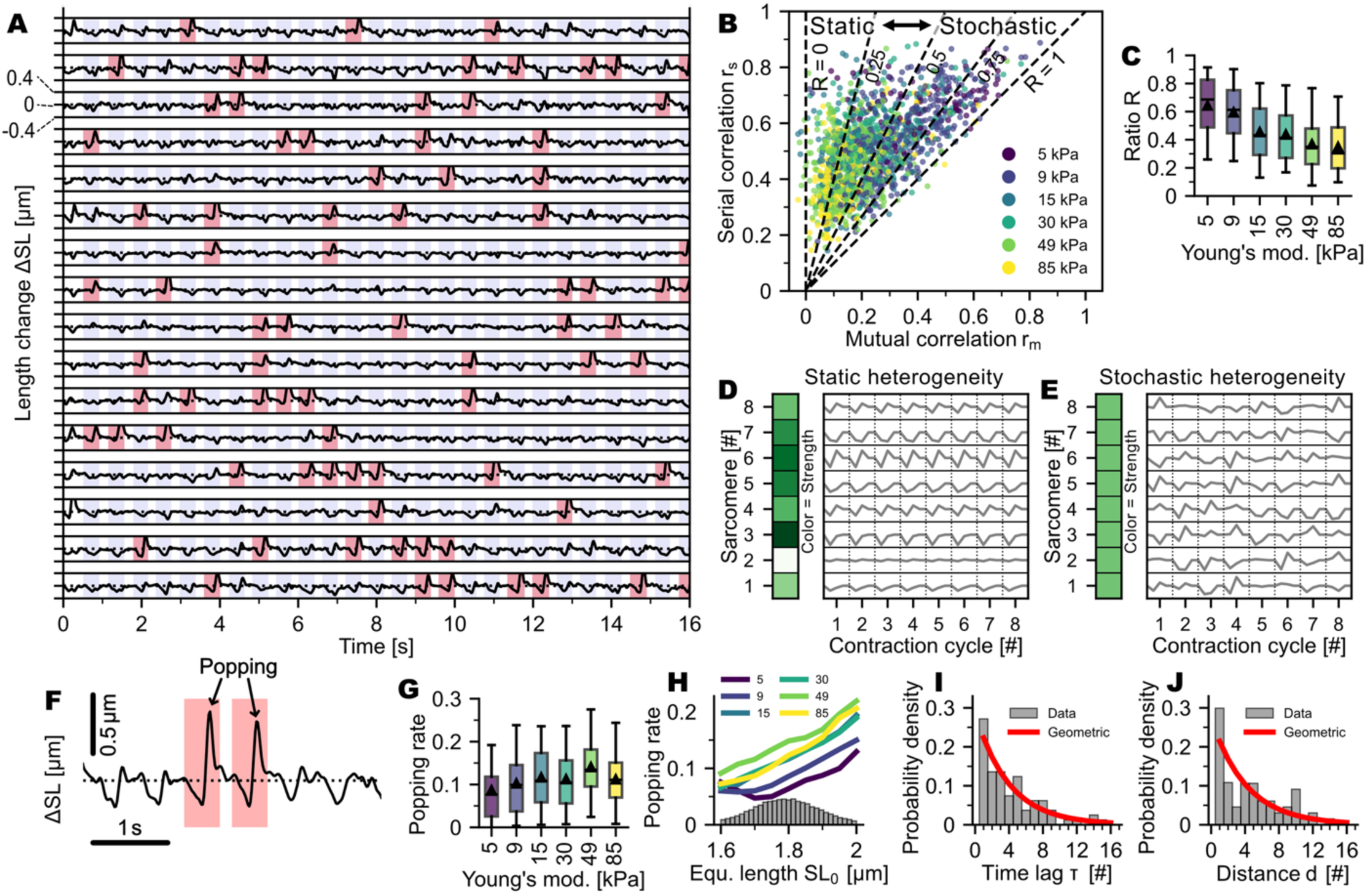
Static versus stochastic heterogeneity in the popping dynamics. **(A)** Representative time-series of sarcomere length changes ΔSL of a myofibril in one representative cardiomyocyte on a 30 kPa substrate. Popping events, defined as sarcomere elongations beyond 0.25 μm within one contraction cycle, are marked in red. Contraction intervals are marked with a blue background. **(B)** Correlation analysis showing mutual correlation (𝑟*_m_*) versus serial correlation (𝑟*_s_*). The x-axis shows the mutual correlation of motion between different sarcomeres (i ≠ j), the y-axis shows the serial correlation of different cycles of one sarcomere (i = j, k ≠ l). Dashed lines delineate regions of static heterogeneity (left) and stochastic heterogeneity (right). Data points are colored by substrate stiffness (5-85 kPa). **(C)** Ratio R between average mutual and serial correlation of ΔSL, serving as a measure for the degree of stochasticity in motions, for different substrate stiffnesses. **(D,E)** Illustrative computer-generated sketches of sarcomere length changes ΔSL in different contraction cycles for purely static heterogeneity (D) and purely stochastic heterogeneity (E). Color bar denotes sarcomere contractile strength. **(F)** Zoomed view of consecutive experimentally observed contraction cycles showing popping events (red shaded regions) where sarcomeres elongated beyond the threshold of 0.25 µm during contraction. **(G)** Overall popping frequencies (in units of 1/contraction cycle) for different substrate conditions. **(H)** Popping frequency as a function of sarcomere equilibrium length SL₀. Lines show averages for different substrate stiffnesses, with underlying distribution of equilibrium lengths shown below in gray. **(I,J)** Probability density distributions of time lag (I) and distance (J) between popping events for single LOI compared with corresponding geometric distributions (red lines). **Data and statistics:** Panels B,C,G-J show data from 2,321 LOIs (5 kPa: 184, 9 kPa: 580, 15 kPa: 392, 29 kPa: 319, 49 kPa: 503, 85 kPa: 343). Box plots show quartiles, mean (triangle) and median (line), with whiskers representing the 5th and 95th percentiles. Statistical analysis was performed using Kruskal-Wallis and Dunn’s post hoc tests, with significance set at p < 0.05. All differences were significant.

where *t* represents the discrete time index (frame number) within a contraction cycle, *T* is the total number of frames capturing one contraction cycle, and 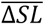 denotes the mean value of the sarcomere length change over the cycle.

For each LOI, we calculated two correlation measures. The mutual correlation coefficient 𝑟*_m_* measures the synchrony between sarcomeres within the same contraction cycle, calculated as 𝑟*_m_* = 〈𝑟(𝑖, 𝑗, 𝑘, 𝑘)〉*_i_*_≠*j*_, where the averaging is performed over all pairs of different sarcomeres (𝑖 ≠ 𝑗) within the same contraction cycle 𝑘. The serial correlation coefficient 𝑟*_s_* quantifies the consistency of individual sarcomeres across different contraction cycles, calculated as 𝑟*_s_* = 〈𝑟(𝑖, 𝑖, 𝑘, 𝑙)〉*_k_*_≠*l*_, where the averaging is performed over all pairs of different contraction cycles (𝑘 ≠ 𝑙) for the same sarcomere 𝑖.

The serial and mutual correlations both were maximal on 5 kPa substrates and declined steadily with increasing substrate stiffness. The decrease of mutual correlation reflects the decrease of synchrony between sarcomeres due to the tug-of-war competition imposed by the rigid constraints on the myofibril level (**Fig. 3B**). The decrease of serial correlation indicates an increased beat-to-beat variability of the motions of individual sarcomeres (**Fig. 3B**). Interestingly, the mutual correlation declined more strongly than the serial correlation, by 50% to more than 90% from 5 - 85 kPa.

We introduce the ratio 𝑅 = 𝑟*_m_*⁄𝑟*_s_* of serial to mutual correlation coefficients to distinguish static from stochastic heterogeneity (**Fig. 3B,C**). When 𝑅 = 1, the heterogeneity is completely stochastic. Random shuffling of sarcomeres (*i, j*) and contraction cycles (𝑘, 𝑙) would in that case not affect *R*. When 𝑅 ≪ 1, the heterogeneity is largely static. In this case, mutual correlations are much smaller than serial correlations, indicating that the motion varies much more between sarcomeres than for a given sarcomere from one contraction cycle to another (**Fig. 3D,E**). The broad distribution of *R* values for all examined LOIs shows that the heterogeneity was neither fully stochastic nor fully static but rather distributed across the full spectrum from stochastic to static (**Fig. 3C**). We examined *R* for different substrate conditions and found a strong dependence on substrate elasticity for both length change and velocity correlations (**Fig. 3B**). Heterogeneity was mostly stochastic for 5 kPa and 9 kPa substrates and increasingly static for stiffer substrates with non-physiological elasticities.

### Sarcomere popping events are stochastically independent and not only occur in structurally “weak” sarcomeres

Beyond the heterogeneous contraction patterns, we observed discrete mechanical instabilities in the form of sudden sarcomere elongations, mostly at the end of contractions, which we termed “popping” events. Defined as sarcomere elongations exceeding 0.25 μm (**Fig. 3A,F**), these events likely represent passive yielding where threshold tension triggers an avalanche-like release of myosin heads while neighboring sarcomeres continue to contract or relax in a more direct manner without overshoot.

The rate of popping events showed a clear dependence on substrate stiffness (**Fig. 3G**). Stiffer substrates (49-85 kPa) led to higher popping rates than softer substrates (5-15 kPa), with median frequencies increasing from approximately 0.08 events per sarcomere per cycle on soft substrates to over 0.14 on the stiffest substrates. Importantly, even on the softest substrates, popping frequencies remained finite, indicating that sarcomere popping is an intrinsic feature rather than a pathological response to specific mechanical conditions. This baseline level of popping across all substrate stiffnesses demonstrates that mechanical instabilities are fundamental to sarcomere dynamics.

Popping frequency also correlated with sarcomere equilibrium length (𝑆𝐿₀), with longer sarcomeres exhibiting higher popping rates across all substrate conditions (**Fig. 3H**). However, the distribution of equilibrium lengths (gray histogram in **Fig. 3H**) was relatively tightly centered around 1.8 μm. Crucially, the popping rate remained consistently non-zero also across the entire range of equilibrium lengths, even in LOIs where length variability was minimal. This demonstrates that while resting length heterogeneity contributes to popping behavior, it is not the primary cause of these mechanical instabilities.

To assess the stochastic nature of popping events, we analyzed the distributions of time intervals and spatial distances between consecutive popping events. We assessed the distributions of distance (𝑑) and time gaps (𝜏) for all LOIs and compared them with geometric distributions 𝐺(𝑘) with respective event probabilities (**Fig. 3I,J**). Using the Kolmogorov-Smirnov test, we could not reject the hypothesis that popping events are stochastically independent for 49% of LOIs with respect to distance and 47% of LOIs with respect to time gaps (p-value = 0.01). This substantial proportion of LOIs showing stochastically independent popping events demonstrates that popping has a stochastic component, independent of deterministic factors such as sarcomere strength heterogeneity or structural variations. While the remaining LOIs show non-random clustering patterns—likely reflecting the influence of local mechanical coupling or morphological heterogeneities—the underlying stochastic mechanism appears to be intrinsic to sarcomere mechanics itself.

### A mesoscopic Langevin framework for coupled sarcomere dynamics

To rationalize the observed complex sarcomere dynamics, we developed a model (**Fig. 4A**) describing the dynamics of each sarcomere 𝑖 by an underdamped Langevin equation—a second-order stochastic differential equation. We represent a sarcomere mechanically as composed of three parallel components: an active force generator, contributing force 𝐹*_a_*_,*i*_, a viscous damper, contributing force 𝐹*_d,i_*, and an elastic element, contributing force 𝐹*_s,i_*. Since we neglect physical mass, forces at all nodes between sarcomeres are instantaneously balanced, meaning every sarcomere experiences the same external load 𝐹*_m_*. However, this external load is not instantaneously balanced by the internal forces (𝐹*_a_*, 𝐹*_d_*, 𝐹*_s_*) within each sarcomere due to the transient nature of the contraction. The resulting force imbalance distributes differently among the three elements from sarcomere to sarcomere, driving asynchronous dynamics (**Fig. 4A**). To capture this non-equilibrium behavior, we introduce the effective parameter µ, which acts as a quasi-inertial mass. The equation of motion reads:

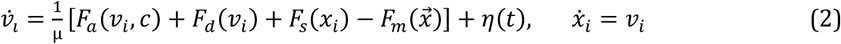

**Figure 4:**
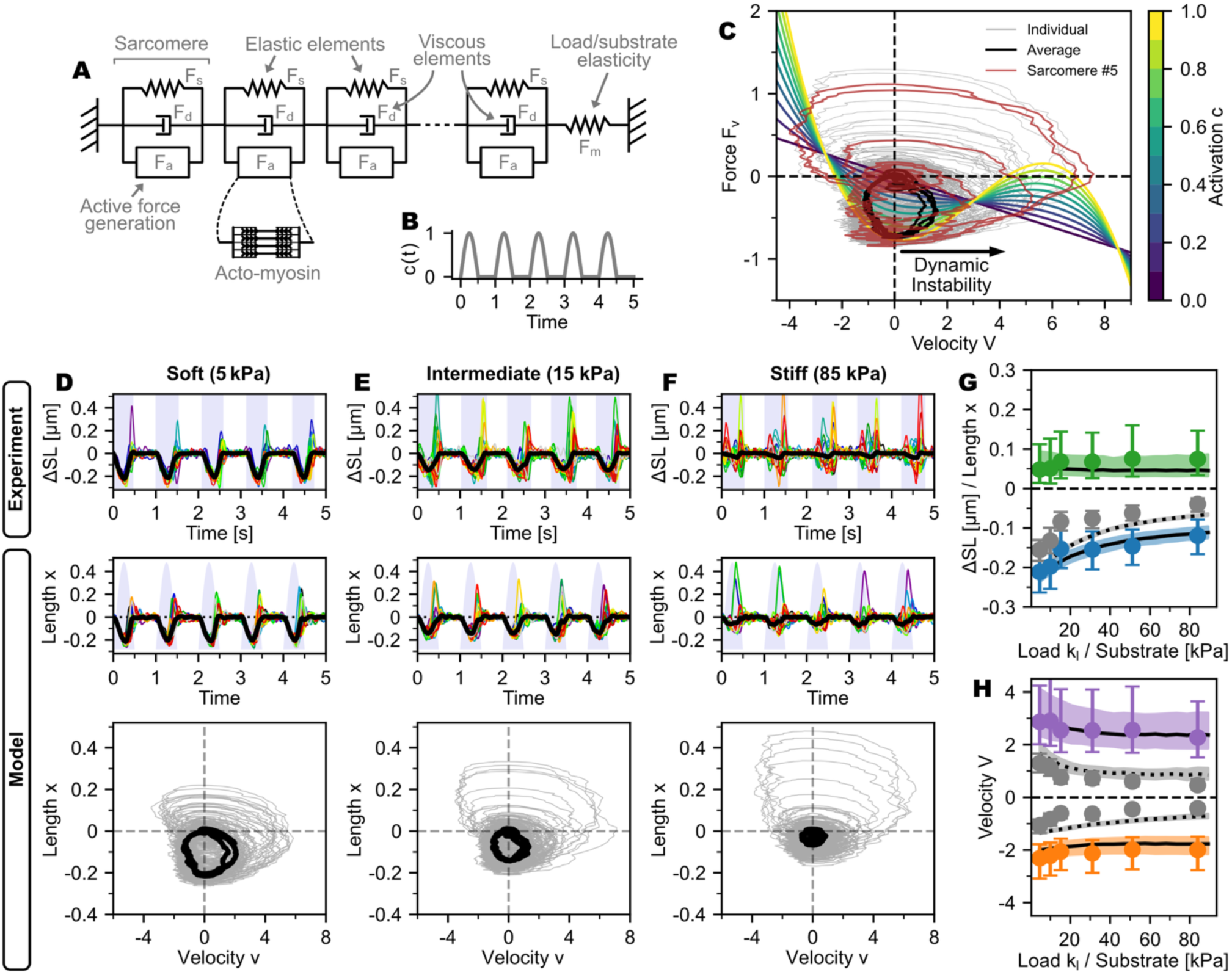
Mesoscopic model of coupled sarcomeres with non-monotonic force-velocity dynamics and comparison with data. (**A**) Schematic of the myofibril model where each sarcomere comprises three parallel components: an active force generator, creating force 𝐹*_a_*, a viscous element creating force 𝐹*_d_*, and passive elastic element, creating force 𝐹*_s_*. Sarcomeres are mechanically coupled in series; such that an external force 𝐹*_m_* generated by elastic substrate deformation is transmitted uniformly through all sarcomeres in the chain. (**B**) Activation imposes a time-dependent modulation of the active force throughout the contraction cycles, with a waveform 𝑐(𝑡). (**C**) Smooth curves: Activation-dependent force-velocity curves showing an S-shaped non-monotonic relationship with two stable branches and an unstable region with a negative-slope; the force–velocity relations are color-coded by activation level. Superimposed trajectories (gray) are trajectories of 20 individual sarcomeres, generated by Eq. 2. The black trajectory is the myofibril average, and one sarcomere is highlighted in red. (**D–F**) Experimental recordings and corresponding model outputs for cells on soft (5 kPa), intermediate (15 kPa), and stiff (85 kPa) substrates showing timeseries of Δ𝑆𝐿/length 𝑥 with blue-shaded contraction intervals (experiment) or 𝑐(𝑡) (model), together with the associated 𝑣–𝑥 trajectories in phase space; colored traces depict individual sarcomeres, and black traces denote cycle averages. (**G–H**) Extrema (maximum and minimum) of length 𝑥 and velocity 𝑣 per contraction cycle for individual sarcomeres and for the myofibril average as a function of substrate stiffness 𝑘*_l_*; dashed lines indicate the average, solid lines show individual-sarcomere values, colored shaded regions denote the standard deviation, and points with error bars display the experimental data median and inter-quantile range.

where 𝑣*_i_* and 𝑥*_i_* are velocity and length of sarcomere 𝑖, 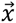 is a vector of all sarcomere lengths, 𝑐(𝑡) is the time-dependent activation, and 𝜂(𝑡) a Gaussian noise term. Multiplying Eq. (2) by µ recovers the form of Newton’s second law (𝐹 = 𝑚 · 𝑎), where 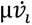 represents the inertial term. While not a physical mass, µ models the characteristic time scale of the system’s kinetic response, reflecting the delayed attachment and detachment rates of actomyosin cross-bridges. This parameter defines how fast the system is drawn to the nullcline determined by the force balance, effectively smoothing instantaneous force fluctuations, representing the system’s persistence against rapid force changes.

This phenomenological underdamped Langevin framework, previously successfully applied to nonlinear systems in general^19^ and to cell migration in particular^20,21^, captures the nonlinear dynamics and emergent collective behaviors observed in our sarcomere system without explicitly modeling the underlying molecular mechanisms. This makes it particularly well-suited for studying the emergence of complex, large-scale behavior from fundamental single-sarcomere interactions.

To model the active actomyosin force 𝐹*_a_*, we chose the 3^rd^ order polynomial ansatz, while the viscous drag 𝐹*_d_* is modeled as a linear friction term:

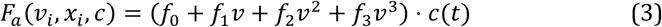

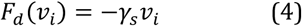

The polynomial in Eq. (3) is a generalist functional form capable of capturing a broad spectrum of force-velocity relationships^21,22^. Multiplying by the time-dependent activation 𝑐(𝑡) makes sure that active force generation scales with calcium availability, reflecting troponin C binding and the resulting availability of myosin binding sites on actin^2^. At high activation (𝑐 ≈ 1), 𝐹*_a_* drives contraction, while at low activation (𝑐 ≈ 0), the viscous drag 𝐹*_d_* dominates, governing passive relaxation dynamics.

In our model, as in previous work^23^, we represent the passive elastic element inside each sarcomere 𝑖 using a piecewise defined function, allowing for different behaviors for shortening and lengthening. We here assume that elastic resistance during shortening is linear (provided by, e.g., microtubules^24^) and quadratic during lengthening (provided by, e.g., titin^23^):

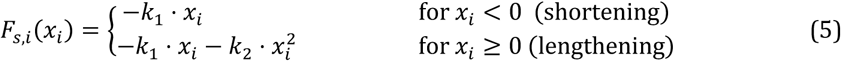

Here, 𝑘_1_ and 𝑘_2_are elastic constants. For simplicity and without loss of generality, we set the sarcomere equilibrium length to zero (𝑥*_eq_* = 0). For serially coupled sarcomeres, the total elastic force on each sarcomere is 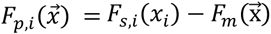 where 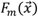 is the external myofibril load. For our experiment of single hiPSC-derived cardiomyocytes on micropatterned elastic soft gels, we assume an auxotonic load 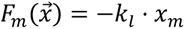 with myofibril length change 𝑥*_m_* = ∑*_j_* 𝑥*_j_* and elasticity 𝑘*_l_*.

We model stochastic fluctuations on the sarcomere level through the random noise term 𝜂(𝑡) = *_t_*(𝑡) + *_a_*(𝑡). This term combines two distinct components: thermal noise 𝜂*_t_*(𝑡) and active noise 𝜂*_a_*(𝑡), originating from the stochastic dynamics of the collective action of the myosin motors in the sarcomere. As shown previously^25^, these fluctuations arising from the stochastic motor kinetics can be approximated as delta-correlated:

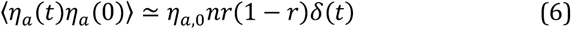

with system-specific magnitude 𝜂*_a_*_,0_, the fraction of active motors 𝑛 and the duty ratio of motors 𝑟. These fluctuations originate from the discrete, stochastic switching of individual motors between bound and unbound states^25^. For the active fluctuations, we model 𝑛 with a sigmoidal function with steepness 𝛽 dependent on velocity 𝑣 dependent on activation 𝑐(𝑡), assuming active shortening and passive elongation of sarcomeres:

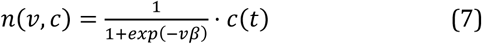

### Data-driven parameter optimization reproduces complex sarcomere dynamics

Our model comprises 10 parameters. To specify the parameters, we simulated 20 serially connected sarcomeres over 10 contraction cycles using the Euler-Maruyama scheme^26^. The cyclic activation was modeled using a simplified custom activation function *c*(*t*) based on a clipped sine with a beating rate matching the experimental data (**Fig. 4B**). Since analytical fitting of such non-linear, stochastic and time-dependent systems to experimental data is impossible, we employed Differential Evolution^27^, a robust gradient-free optimization method, to identify the physical parameters generating the experimentally observed dynamics. The objective was to minimize a loss function based on the Kolmogorov-Smirnov (KS) coefficient, which quantifies the similarity between the distributions of simulated and experimental data on a scale from 0 (identical) to 1 (no overlap).

The model evaluation compared the distributions of sarcomere length changes and velocities from simulations with representative experimental LOIs from substrates (5, 15, and 85 kPa, mapped to *kl* = 0.5, 1.5 and 8.5 in our 1-D model; 𝑘*_l_* is unitless, so only the ratios between values are meaningful — rescaling 𝑘*_l_* leaves model output unchanged under correspondingly rescaled parameters) covering the full range of mechanical loads. To account for the transient activation dependency, we divided the trajectories into four distinct phases: quiescent, early activation, mid-activation, and late activation, and compared the distributions for each phase using the KS metric. To match the experimental temporal resolution, model data was subsampled to 16 ms frame intervals. For each parameter set during optimization, all three substrate conditions were simulated, and the KS coefficient was computed for each condition and phase. The final loss was then calculated as the mean loss over all three conditions and four contraction phases, yielding a single metric quantifying the overall model-experiment agreement. The 10 model parameters were optimized within prespecified bounds (**Table S2**).

This optimization process yielded a model in excellent agreement with the experimental data (**Fig. 4D-F**). The average KS coefficient across all conditions and phases was ∼0.05, confirming a high degree of similarity between the simulated and measured distributions of individual sarcomere lengths. Ultimately, we inferred a single, unified model capable of replicating dynamics across all substrate elasticities by adjusting only the substrate stiffness parameter. The model correctly captures the extremal trends (minima and maxima) of decreasing sarcomere displacement and velocity with increasing substrate stiffness at both the individual and average sarcomere levels (**Fig. 4G, H**). While the model predicts slightly stronger average contraction and velocity extrema than measured, the stiffness dependence of these extrema is consistent with experimental observations (gray data in **Fig. 4G, H**). Note that the model correctly replicates the emergence of smooth and periodic contraction at the myofibril level from the highly stochastic and heterogeneous dynamics of individual sarcomeres (**Fig. 4D-F**).

### Non-monotonic force–velocity dynamics drive tug-of-war and limit cycles in coupled sarcomeres

Our data-driven inference procedure reveals, without making this an *a priori* assumption, that the active force 𝐹*_a_* exhibits a non-monotonic force-velocity relationship (**Fig. 4C, Movie S2**), with branches separated by an unstable region of negative slope (𝑑𝐹/𝑑𝑣 < 0), corresponding to negative friction. This relationship is dynamically modulated by the activation 𝑐(𝑡), creating a complex time-dependent force landscape that governs sarcomere behavior.

The model is effectively analogous to a Van-der-Pol relaxation oscillator in force-velocity space^19^. As total myofibril shortening proceeds, the external mechanical load from the substrate increases, slowing down contraction velocities of all sarcomeres. The non-monotonic force-velocity relationship implies that there is an intrinsic threshold force at which sarcomeres become dynamically unstable. Physically, this corresponds to a tipping point where the stochastic unbinding of a few myosin cross-bridges triggers an avalanche-like release of the remaining more strongly loaded cross-bridges, switching the sarcomere from contraction to rapid elongation.

This instability has different consequences in the different phases of the contraction cycle: During the initial contraction phase, joint sarcomere shortening drives the velocity toward the instability threshold. Through dynamic selection, stochastic fluctuations (modeled by 𝜂) then push random sarcomeres past the tipping point. Because activation remains high, this elongation is transient; as a particular sarcomere lengthens, load quickly redistributes to its passive elements and while other sarcomeres take up the slack, it ends up back on the contracting branch. This rapid switching generates the stable limit-cycle high-frequency oscillations observed in individual sarcomeres (**Fig. 2H-J**).

During relaxation (declining activation), the instability threshold decreases (**Fig. 4C**). On stiff substrates, where external load remains high, this lower threshold causes a larger fraction of sarcomeres to slip past the instability point. Unlike in the first regime, the declining active force allows for large, sustained “popping” excursions (**Fig. 4D-F**). We observed that only a subset of sarcomeres pop (∼10%), which effectively buffers the load for the remaining sarcomeres, allowing them to relax smoothly without over-extension.

Stochastic fluctuations 𝜂(𝑡) (Eq. 6) are essential to reproduce the heterogeneous sarcomere behavior seen experimentally, including beat-to-beat variability, competition, and popping events. Removing the stochastic drive from the equation of motion (Eq. 2) in simulations collapses trajectories, inconsistent with experiments (**Fig. 5A,B**). To probe robustness against fixed force non-uniformities, we introduced static variability by scaling each sarcomere’s active force 𝐹*_a_* with a constant, factor drawn for each sarcomere in the chain of 20 from a distribution with standard deviation 𝜎*_a_*. Functionally, this assigns each sarcomere a slightly different maximum force threshold. This force variability alone biases trajectories and creates deterministic static heterogeneity during contraction according to each sarcomere’s strength (**Fig. 5C**), also inconsistent with our data. Adding stochastic fluctuations leads to trajectories most closely resembling the experimental data (**Fig. 5D**).

**Figure 5:**
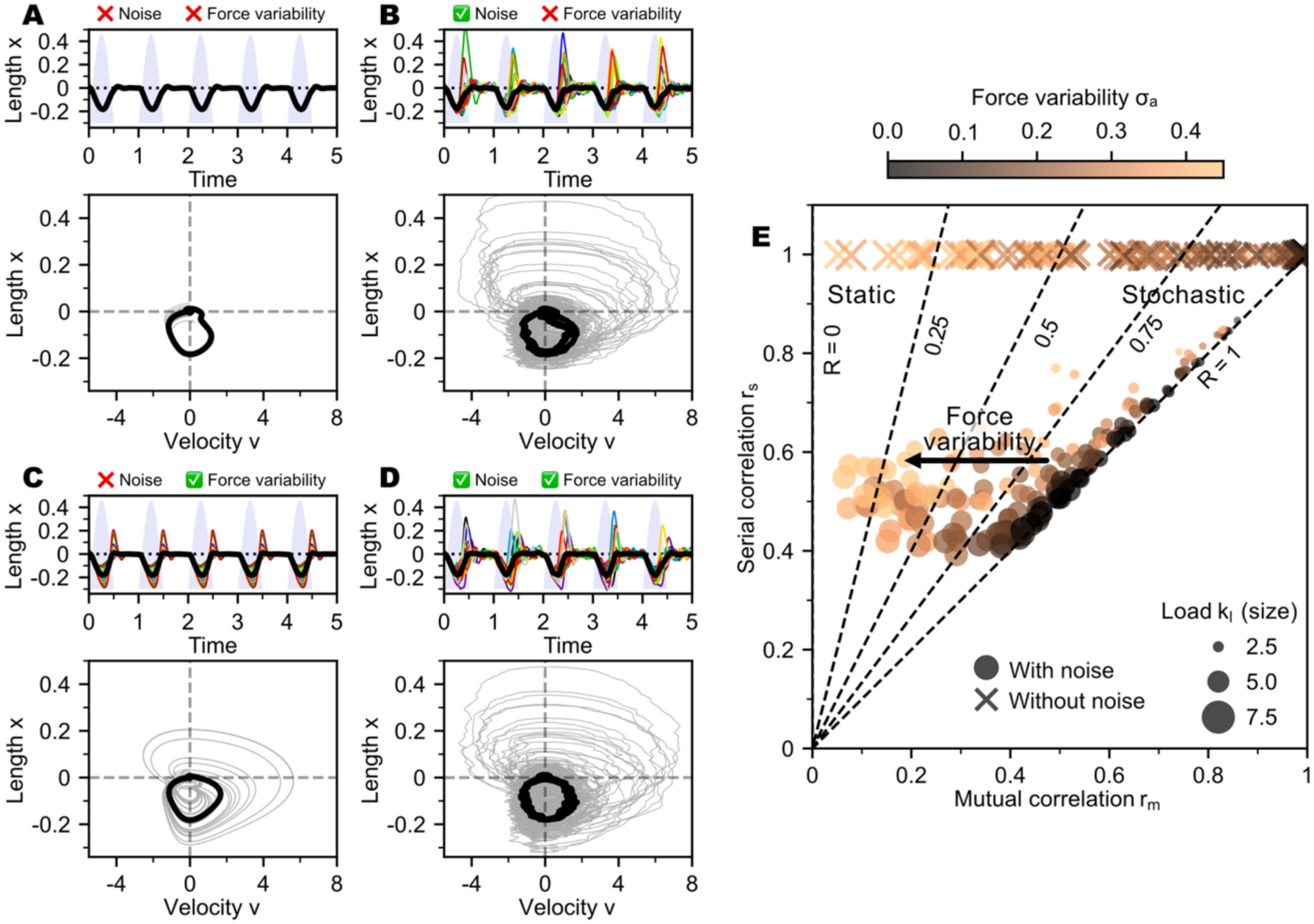
Simulation of static and stochastic heterogeneity in sarcomere dynamics. (**A–D**) Top panels: overlaid timeseries of individual sarcomere length changes (colored curves) and the ensemble average (thick black curve). Bottom panels: trajectories in phase space (length change vs. velocity) of individual sarcomeres (gray) and ensemble average (black). **(A)** Uniform ensemble (no noise, no force variability). **(B)** Stochastic heterogeneity driven by noise only. **(C)** Static heterogeneity driven by intrinsic force variability (𝜎*_a_*) only. **(D)** Mixed, static and stochastic heterogeneity with both noise and force variability. (**E**) Serial correlation 𝑟*_s_* versus mutual correlation 𝑟*_m_* across load levels and force variability; marker size encodes the dimensionless simulated substrate stiffness 𝑘*_l_*, and color shade indicates the standard deviation of the multiplicative factor modulating each sarcomere’s active force. Dashed lines mark the transition from purely static heterogeneity (𝑅 = 0) to purely stochastic heterogeneity (𝑅 = 1). Circles denote simulations with noise/stochastic fluctuations, and crosses denote deterministic simulations without noise/stochastic fluctuations.

Correlation analysis (**Fig 5E**; compare with experimental data in **Fig. 3B**) of the simulated data with different levels of static heterogeneity 𝜎*_a_* shows, as expected, that without stochastic fluctuations, the serial correlation of sarcomeres is always 1, meaning that individual sarcomeres always behave the same way through contraction cycles. In contrast, simulations with stochastic fluctuations maintain stochastic heterogeneity (R > 0.8) even with substantial static force heterogeneity (up to 𝜎*_a_* ≈ 0.2) (**Fig. 5E**).

## Discussion

We combined high-speed imaging of human induced pluripotent stem cell (iPSC)-derived cardiomyocytes with detailed AI-based quantitative analysis. We discovered rich subcellular dynamics that remain hidden at the whole-cell level. While overall cell contraction cycles were regular, individual sarcomeres displayed rapid oscillations, reversible popping events, and stochastic beat-to-beat variability within the same myofibril. Increased substrate stiffness reduced overall cell contraction amplitudes, but had only a weak effect on single-sarcomere length-changes and contraction velocities. Rigid substrates instead promoted tug-of-war-like interactions and loss of synchrony between sarcomeres along myofibrils.

To interpret these observations mechanistically, we developed a data-driven, mesoscopic modeling framework describing transient, non-equilibrium dynamics and mechanical coupling among serially connected sarcomeres. The model relies on two components that accurately reproduce the experimentally observed dynamics: a non-monotonic force-velocity relationship that generates a dynamic instability at a critical force and intrinsic stochastic force fluctuations at the sarcomere level. This approach differs from classical modeling of skeletal muscle based on a force-length relationship. We find that our model is more appropriate than such models for cardiomyocytes that operate in a regime of rapid transient activation-relaxation cycles, viscous drag, and velocity-dependent force generation.

The dynamic instability central to our framework arises from the non-monotonic shape of the force-velocity relationship. Note that this functional form was not an initial assumption but that it emerged when optimizing the parameters of the very general modeling ansatz to fit our experimental data. This result connects to foundational, reductionist theories of coupled molecular motors working against an external load^28–31^, which predict that at a critical load, a collective, avalanche-like unbinding of motors from their track (e.g. myosin heads from actin in a sarcomere) induces a dynamic phase transition from active forward motion against the load to rapid passive slipping. Such instabilities have so far only been reported in reconstituted motility assays^30,32^. We have here observed an analogous instability in cardiomyocytes manifesting itself in the observed switching between contraction and expansion and the popping events of individual sarcomeres. If sarcomeres followed a monotonic force-velocity relationship, increasing load would simply cause uniform slowing or stalling. Small non-uniformities caused by fluctuations would decay, returning the system to a uniform state. The non-monotonic force-velocity relationship, instead, amplifies such fluctuations, driving the system into limit cycles that produce the high-frequency oscillations and stochastic heterogeneity we observe (**Fig. 2G**).

What could be the physiological roles of the observed phenomena? The popping events are the most dramatic outcome of the instability-amplified stochastic noise. Their reversible nature, ubiquity across all tested conditions, and prevalence at the end of contractions – where declining activation reduces the fraction of attached cross-bridges – are all consistent with a dynamic instability rather than structural failure. We therefore view popping as an intrinsic outcome of transient, non-equilibrium sarcomere dynamics. While possibly similar popping has been described in skeletal muscle^33^, particularly in the context of residual force enhancement^34^, the observation in rhythmically contracting cardiomyocytes is a key new finding of this study. A physiological function of popping might be that intermittent yielding in a subset of sarcomeres accelerates myofibril-level relaxation toward the onset of the next diastole, helping tension decay even while individual sarcomere lengths remain non-uniform. A physiological role of stochastic heterogeneity of yielding might be the limitation of sarcomere damage. Our data indicate a mixture of static and stochastic heterogeneity. Some sarcomeres exhibit persistent differences suggestive of structural or compositional non-uniformities, yet many myofibrils show strong, cycle-to-cycle stochastic variability, including popping. As our simulations demonstrate, even in the presence of low to moderate sarcomere strength variability, the interplay between stochastic fluctuations and the dynamic instability prevents deterministic yielding. The interplay between intrinsic fluctuations and the dynamic instability randomizes yielding events across cycles and sarcomeres, preventing the repetitive overload of any single element and reducing the likelihood of localized damage.

Stochastic heterogeneity thus provides a robust mechanism for maintaining structural and functional homeostasis. Healthy adult cardiomyocytes—with their more ordered myofibril and sarcomere architecture compared with stem-cell-derived cardiomyocytes^35^—are expected to show smaller static non-uniformities while still exhibiting stochastic heterogeneity. While speculative for human cardiomyocytes, this hypothesis is supported by in-situ measurements of single-sarcomere dynamics in mouse myocardium, which found stochastic heterogeneity^6^.

In diseased and disordered myofibrils, static non-uniformities may overwhelm the protective stochastic mechanism ^14,35,36^ and instability may then be consistently triggered at the most vulnerable sarcomeres, concentrating mechanical stress and leading to fatigue and maladaptive remodeling. This perspective provides a more nuanced view than those based on fixed “weak–strong” contrasts^14^, suggesting that pathology may arise from local structural weakness, but that that weakness would have to overwhelm the protective load-sharing mechanism. The phenomenon that amplified random noise can enhance responsiveness and stabilize dynamics has also been described in systems showing stochastic resonance^37^.

The presented framework is mesoscopic by design, omitting explicit molecular details such as calcium kinetics or individual cross-bridge states. Instead, we adopted a top-down approach, inferring the emergent mesoscopic physics directly from the experimental data. We utilized a mesoscopic Langevin framework in the hydrodynamic limit, where the deterministic force term represents the collective average of the underlying molecular kinetics and mechanical interactions, while the stochastic term accounts for unresolved microscopic fluctuations, following the general principles of active matter theory applied to actomyosin systems^38,39^. The strength of this coarse-grained approach is that it reveals emergent physical principles—such as the non-monotonic force-velocity relationship—that would remain obscured in parameter-heavy molecular models. The model’s predictive power, with only 10 parameters capturing dynamics across three substrate stiffnesses and four contraction phases, validates this strategy.

Our combined experimental and modeling platform offers a powerful tool to dissect the specific molecular drivers of collective sarcomere dynamics in future studies. By applying pharmacological perturbations or specific mutations, one could link distinct molecular defects — such as altered cross-bridge kinetics, calcium sensitivity, or titin stiffness — to quantitative changes in the model’s physical parameters. This would establish a rigorous mapping between molecular function and myofibril stability, enabling a systematic analysis of how specific pathologies disrupt the contractile homeostasis of the heart.

## Materials and Methods

### Generation and culturing of hiPSC ACTN2-Citrine-derived cardiomyocytes

Cardiomyocyte differentiation of a ACTN2-Citrine reporter hiPSC line^40^ was performed according to Tiburcy *et al.*^41^. hiPSC-ACTN2-Citrine-derived cardiomyocytes were cultured in 6 well plates (Cat 3516, Corning) in serum-free “cardio” medium (0.4 mM Ca^2+^, RPMI 1640 with GlutaMAX (Cat 61870, Invitrogen), 1% Penicillin/Streptomycin (Cat 15140, Invitrogen), 2% B27 supplement (Cat 17504-044, Invitrogen) at 37 °C in a 5% CO2 incubator with culture medium changes at every other day. For re-seeding, cells were detached using Accutase^®^ digestion medium (StemPro^®^ Accutase^®^ cell dissociation reagent (Cat A11105-01, Gibco), 0.025% Trypsin (Cat 15090-046, Gibco), 20 µg/mL DNaseI (Cat 260913, Calbiochem) for 15-20 min at 37°C. Digestion was stopped using “cardio” medium supplemented with 5 µmol/L Rock Inhibitor (Stemolecule Y27632, Cat 04-0012-10, Reprocell) three times the volume of the Accutase^®^ mix. Cell clumps were separated using a 100 µm cell strainer. Cells were seeded using ∼150,000 cells per micropatterned patterned substrate of ∼1 cm^2^ size and cultured 24 hours in “cardio” medium with 5 µmol/L Rock Inhibitor. Cardiomyocytes on soft gels were maintained in serum-free “cardio” medium with daily medium changes at 37 °C in a 5% CO2 incubator for up to 30 days. Cardiomyocytes were imaged after a maturation period of 20-30 days post seeding on the soft gels.

### Sample preparation and measurement

For live-cell video imaging, the soft substrates were mounted in a custom-built holder for round Ø25 mm glass cover slides with serum-free medium. Movies of beating cardiomyocytes were obtained with a confocal microscope (TCS SP5 II, Leica, Germany) at 37°C and 5% CO2. We used an 8,000 Hz resonant scanner with bidirectional scanning mode and recorded up to 20-30 s long movies of 1024 × 200 pixels at 67 frames per second, i.e. a temporal resolution of 15 ms.

### Fabrication of cell-adhesive micropatterns on polyacrylamide soft gels

Silicon wafers (Microchemicals GmbH, Ulm, Germany) were coated with SU-8 photoresist (Series 3005, MicroChem, Newton, USA) using a two-step spin-coating process. The wafers were then exposed to UV light through custom photomasks (Compugraphics, Jena, Germany) and developed to create photoresist masters (details in Ref^40^). Polydimethylsiloxane (PDMS) stamps were produced by mixing PDMS and curing agent 10:1 (Sylgard 184 kit, Dow Corning) and pouring it onto the photoresist masters. After degassing and curing, the PDMS was cut and peeled off to create the stamps. Micropatterned polyacrylamide gels were prepared by treating PDMS stamps with plasma to make them hydrophilic, then incubating them with 0.1 mg/ml Synthemax™ (Cat 3535, Corning). The protein-coated stamps were placed on plasma-cleaned glass coverslips and weighted to transfer the protein. Gel solutions of acrylamide and bis-acrylamide were prepared to achieve different elastic moduli (**Table S1**), measured with a rheometer (Physica MCR 501 Rheometer, Anton Paar, Austria). The gel solution was polymerized with the Synthemax-patterned glass on top. The gels were stored in PBS, then washed before use.

### Microscopy data analysis and statistical analysis

For processing of the live-cell confocal movies, and tracking and analysis of sarcomere trajectories, our Python package SarcAsM (Sarcomere Analysis Multitool) was used^15^. All additional analyses and illustrations were created using custom scripts written in Python. All data is displayed as mean ± standard deviation unless indicated otherwise. Whenever applicable, we employed the Kruskal-Wallis test, a non-parametric method, for hypothesis testing, and proceeded to a post-hoc multi-comparison using Dunn’s test, setting the threshold for statistical significance at a p-value of less than 0.01. All condition differences were deemed significant except where explicitly marked as not significant (n.s.).

### Computational model

We implemented the mesoscopic model of coupled sarcomeres in Python 3.12, making use of the NumPy^42^ and SciPy^43^ libraries. The model describes sarcomere dynamics through serially coupled underdamped Langevin equations, which is an approach suited to capturing emergent collective behaviors at the myofibril scale without explicitly modelling molecular-level details. We solved the system of stochastic differential equations numerically using the Euler-Maruyama scheme^26^ with a time step of 2 ms. The dimensionless load parameter 𝑘_0_was provided as a simulation input, defined for simplicity as 0.1 times the substrate elasticity (in kPa). The remaining model parameters were estimated using Differential Evolution^27^, a global optimization algorithm, to minimize the Kolmogorov–Smirnov distance between simulated and experimental distributions.

## Data and Code Availability

All data, analysis scripts, and source code supporting this study are publicly available. The primary software packages developed for this research, SarcAsM and the computational model, are available on GitHub at https://github.com/danihae/SarcAsM and https://github.com/danihae/SarcomereModel, respectively. The complete experimental dataset, alongside code to reproduce the analyses in this manuscript, is archived on Zenodo (https://doi.org/10.5281/zenodo.17564384). This repository includes a representative dataset of 25 raw microscopy movies with their corresponding processed line-of-interest (LOI) sarcomere trajectories for each of the six substrate stiffness conditions, as well as the aggregated data frame used to generate all figures.

## Acknowledgements

We thank Florian Rehfeldt, Andrei Vilfan, Lev Truskinowsky, David Brückner, Chase Broedersz and Pierre Ronceray for helpful discussions. We gratefully acknowledge the use of the microscopy facility of the Max Planck Institute for Multidisciplinary Sciences for access to cell culture and high-speed confocal microscopy. CFS and DH would like to thank the Isaac Newton Institute for Mathematical Sciences for support and hospitality during the program “New statistical physics in living matter: non equilibrium states under adaptive control”. DH acknowledges the support from the German Academic Foundation (Studienstiftung des Deutschen Volkes) for providing a doctoral fellowship and the Campus Institute for Data Science (CIDAS) at the University of Göttingen for awarding a postdoctoral fellowship. The research of CFS was supported by the European Research Council under the European Union’s Seventh Framework Program (FP7/2007-2013) with the ERC grant agreement n°340528. WHZ acknowledges the support from the DZHK (German Center for Cardiovascular Research), the German Federal Ministry of Education and Research (IndiHEART; 161L0250A), the German Research Foundation (DFG SFB 1002 C04/S01, IRTG 1816, RTG 2824, EXC 2067-1), and the Fondation Leducq (20CVD04).

## Author contributions

D.H., T.D., W.H.Z. and C.F.S. conceived and designed the study. D.H., L.H., and T.D. developed the methodology. W.H.Z. and C.F.S. supplied the essential resources. D.H., L.H., and T.D. performed micropatterning and cardiomyocyte experiments. D.H. and K.N. analyzed experimental data. D.H. conceived and implemented the theoretical model. D.H. created the visualizations. The original draft was written by D.H., and all authors contributed to the review and editing of the manuscript.

## Materials & Correspondence

For correspondence and requests for materials, please contact Wolfram-Hubertus Zimmermann and Christoph F. Schmidt.

## Supplementary Information

**Figure S1:**
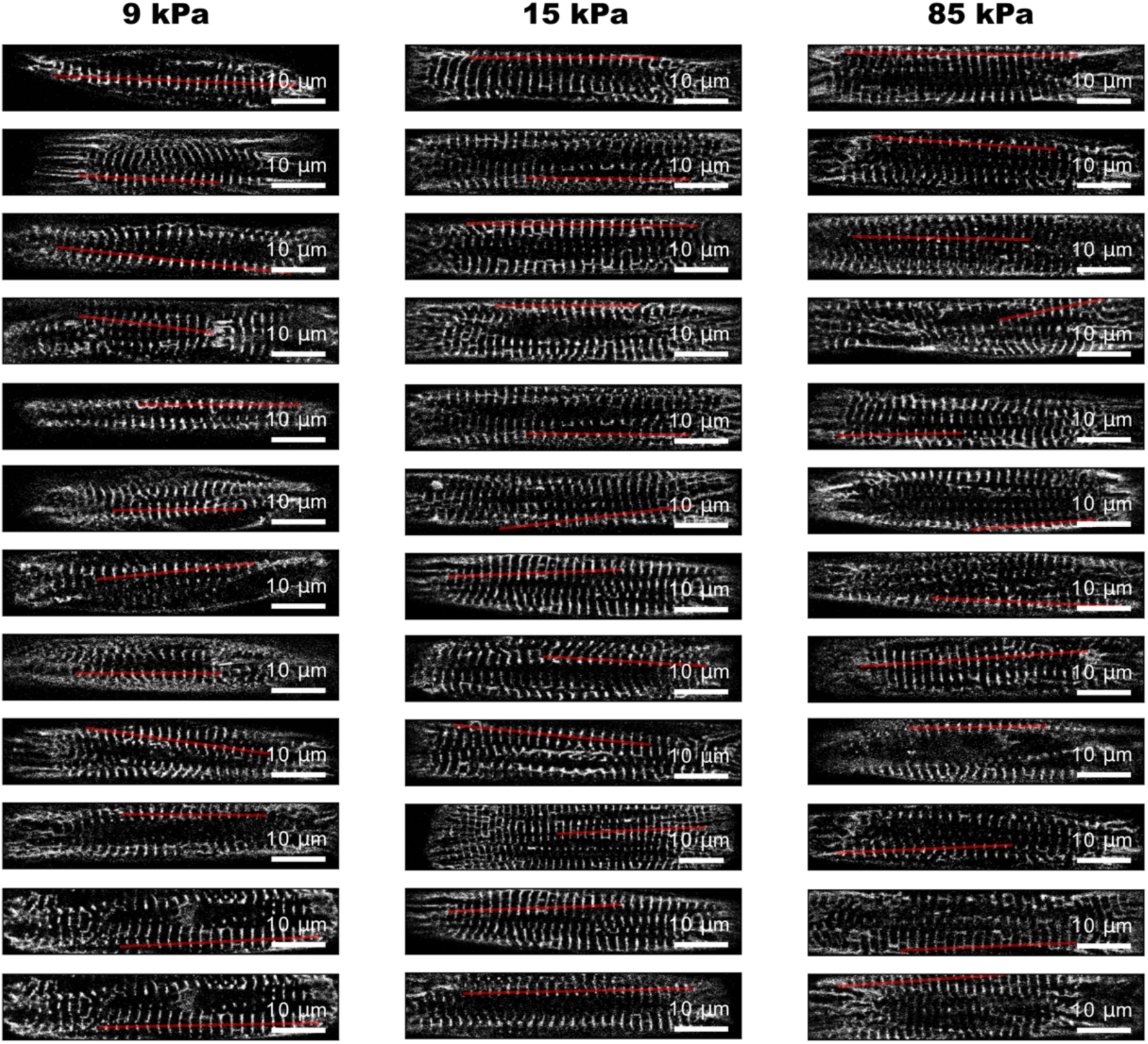
Representative CM selection with automatically and individually detected ROIs. CMs are randomly selected from the data sets for three substrate stiffnesses (9, 15 and 85 kPa); automatically determined ROIs depicted as red lines.

**Figure S2:**
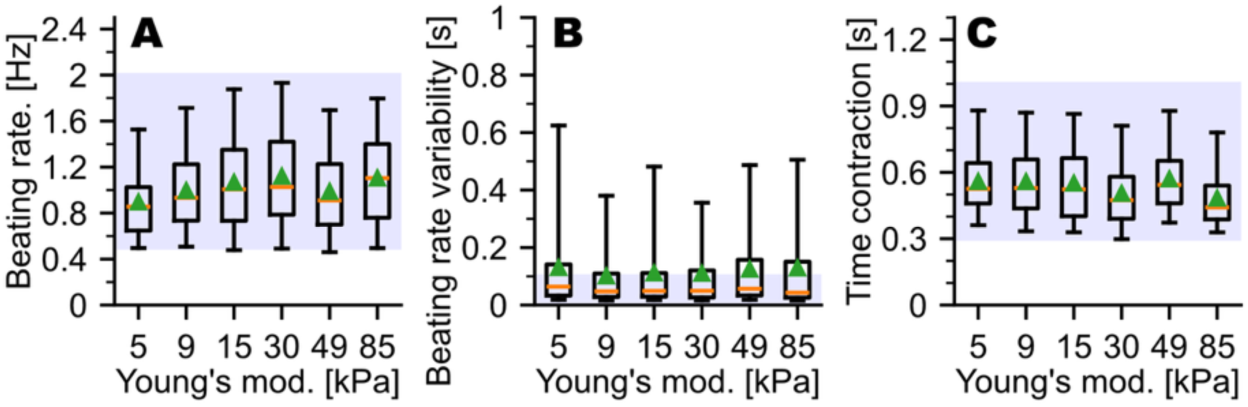
Effects of substrate elasticity on spontaneous beating of cardiomyocytes. (**A**) Spontaneous beating frequencies. (**B**) Beating irregularity: relative standard deviation of beating periods. (**C**) Average length of contraction cycles *Tc*. **A-C** include data from 5,085 LOIs. Boxes show quartiles, red lines the median, green triangles the mean and whiskers the 5th and 95th percentile of the distributions. Blue shaded intervals show included data.

**Table S1:**
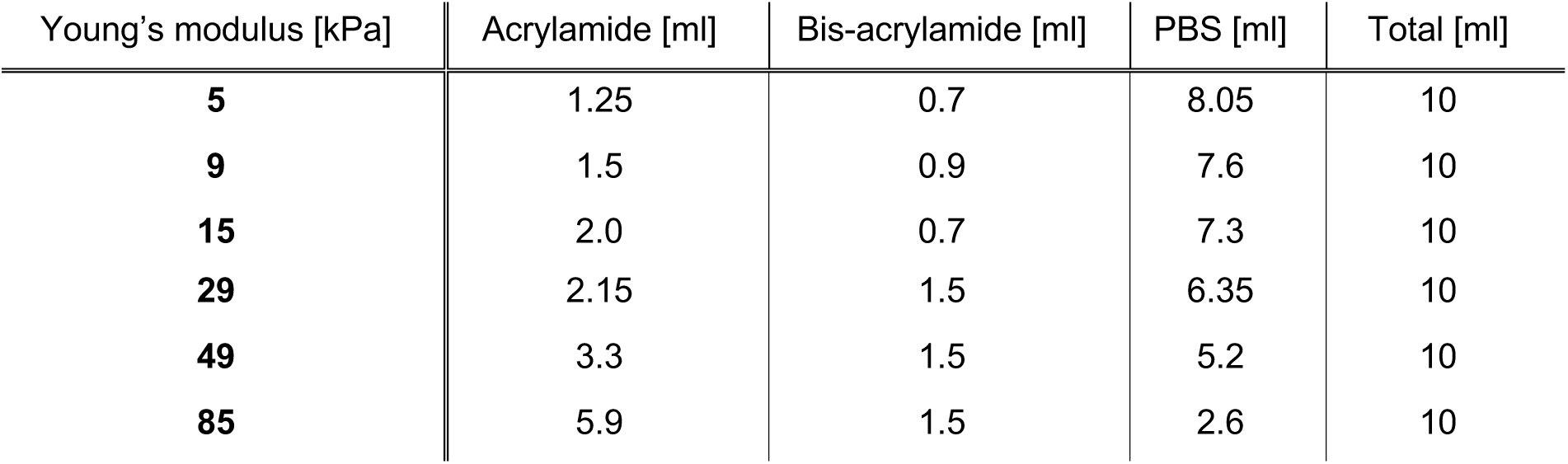
Composition of polyacrylamide soft gels used in the study. All gels used here were made from the same respective stock solution (10 ml). The Young’s modulus was measured for a gel sample from each prepared stock solution using a rheometer (Physica MCR 501 Rheometer, Anton Paar, Germany) using a 25 mm, 2° cone plate with a sample volume of 140 μl. A time sweep (1 h, spacing 30 s, 1% strain, 1 Hz), a frequency sweep (3 measurements per decade, 1% strain, 0.01 – 100 Hz), and an amplitude sweep (3 measurements per decade, 0.01 – 100% strain, 1 Hz) were performed consecutively.

| Young's modulus [kPa] | Acrylamide [ml] | Bis-acrylamide [ml] | PBS [ml] | Total [ml] |
| --- | --- | --- | --- | --- |
| **5** | 1.25 | 0.7 | 8.05 | 10 |
| **9** | 1.5 | 0.9 | 7.6 | 10 |
| **15** | 2.0 | 0.7 | 7.3 | 10 |
| **29** | 2.15 | 1.5 | 6.35 | 10 |
| **49** | 3.3 | 1.5 | 5.2 | 10 |
| **85** | 5.9 | 1.5 | 2.6 | 10 |

**Supplementary Table S2.** Model parameters obtained through differential evolution optimization. Parameters were optimized by minimizing the Kolmogorov-Smirnov statistic between model simulations and experimental data from ACTN2-Citrine hiPSC-derived cardiomyocytes. Search ranges were based on physical constraints and preliminary analyses. Parameter values were rounded.

| Parameter | Symbol | Optimized Value | Search Range | Description |
| --- | --- | --- | --- | --- |
| Inertia | $\mu$ | $6 \times 10^{-3}$ | [0.0001, 0.01] | Pseudo-inertial parameter controlling adaptation timescale to force changes |
| Thermal noise | $\eta_0$ | 2.2 | [0.5, 3.0] | Amplitude of thermal fluctuations (Gaussian noise) |
| Active noise | $\eta_1$ | 9 | [1.0, 10.0] | Amplitude of active fluctuations from stochastic motor dynamics |
| Stiffness (linear) | $k_{\{s,0\}}$ | 2.5 | [0.5, 5.0] | Linear elastic constant for sarcomere shortening and lengthening |
| Stiffness (quadratic) | $k_{\{s,1\}}$ | 3.2 | [0, 10.0] | Quadratic elastic constant for sarcomere lengthening ($x > 0$) |
| Friction | $u_{\{s,0\}}$ | 0.10 | [0.01, 0.2] | Passive friction within sarcomere (velocity-dependent) |
| F-V constant | $f_{\{s,0\}}$ | -0.8 | [-5, 5] | Constant term in polynomial force-velocity relation |
| F-V linear | $f_{\{s,1\}}$ | -0.06 | [-5, 5] | Linear coefficient in polynomial force-velocity relation |
| F-V quadratic | $f_{\{s,2\}}$ | 0.10 | [-5, 5] | Quadratic coefficient in polynomial force-velocity relation |
| F-V cubic | $f_{\{s,3\}}$ | 0.012 | [-5, 5] | Cubic coefficient in polynomial force-velocity relation |

**Movie S1:**
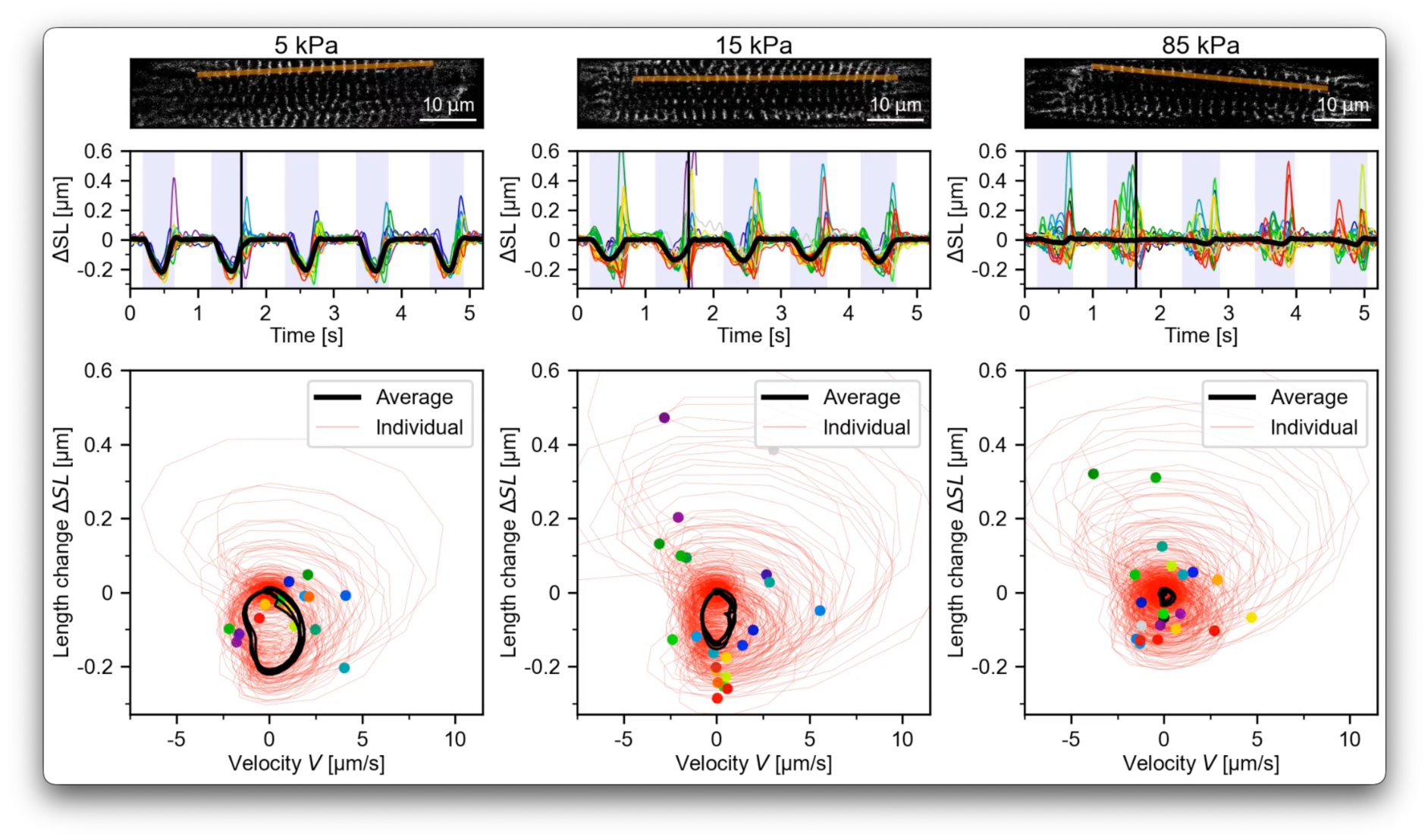
Representative cardiomyocytes showing the effect of substrate stiffness on sarcomere heterogeneity. Real-time confocal movies of three representative ACTN2-Citrine labeled cardiomyocytes (16 ms frame time) on 5 kPa, 15 kPa and 85 kPa substrates. Red lines show lines of interest (LOI) for sarcomere motion tracking and analyses. Middle: Individual sarcomere length changes 𝚫𝑺𝑳_𝒊_(𝒕) (colored lines) and average sarcomere length change 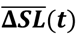 (black line) of myofibril ROIs shown above. Contraction intervals are shown in light blue. Black vertical line indicates time progress synchronized with movies above. Bottom: Length-change velocity phase portraits showing individual sarcomere trajectories (red traces), the average trajectory (black trace), and individual sarcomere positions (colored dots).

**Movie S2:**
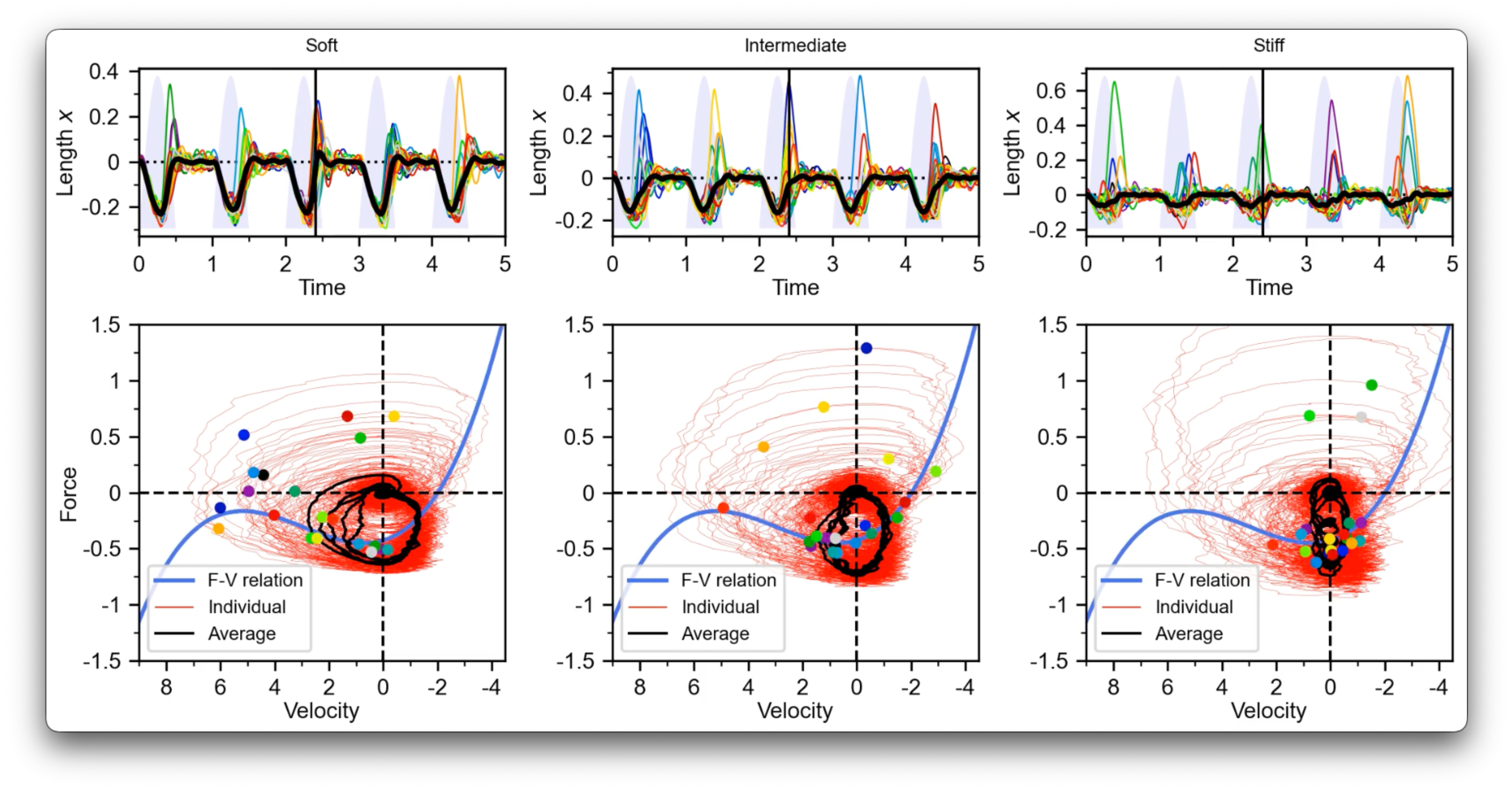
Computational model simulations showing the effect of substrate stiffness on sarcomere coordination. Simulations of sarcomere dynamics under three substrate stiffness conditions (resembling 5 kPa, 15 kPa, and 85 kPa). Top: Individual sarcomere length changes over time (colored traces) and the ensemble average (black trace). The thin vertical line indicates time progress. Bottom: Phase portraits in velocity-force space showing individual sarcomere trajectories (red traces), the ensemble average trajectory (black trace), and instantaneous sarcomeres at each time point (colored dots). The blue curve represents the force-velocity relationship (nullcline).

